# Making friends in an asymmetric game: the establishment of male-female grooming exchanges in vervet monkeys

**DOI:** 10.1101/2025.02.20.639230

**Authors:** Josefien A. Tankink, Maria Granell Ruiz, Erica van de Waal, Carel van Schaik, Redouan Bshary

## Abstract

An important challenge for individuals of group-living species is to build cooperative relationships with new partners. One famous strategy for doing so is “raise-the-stakes”, proposing low initial investments that increase if the partner matches these investments in a series of exchanges. Contrarily, the “all-in” strategy predicts high initial investment that may be maintained, but downregulated if not fully matched. We tested these predictions on grooming exchanges between wild male-female vervet monkeys, a species with regular male dispersal. Contrary to model predictions, we found uneven initial investment between novel partners that reached near reciprocity after approximately six months. To identify which sex altered investment, we examined grooming durations and frequencies of novel and established partners. Females showed consistently high initial investment and gradually reduced grooming over the first six months. Males did not alter grooming duration but had higher grooming frequency after a year of residency. Eventually, grooming exchanges nearly evened out. Female behaviour aligns more closely with the “all-in” strategy, which suggests competition among female groups over dispersing males. Male behaviour partly fits the “raise-the-stakes” hypothesis. The study highlights the complexity of real-life cooperation, emphasizing the need to explicitly incorporate life history parameters to understand cooperative strategies.

## Introduction

Living in stable groups selects for social competence so that individuals can manage cooperation and conflict. One such challenge is to respond to turnover in group composition by building cooperative relationships with new partners. Cooperative behaviour is often prone to exploitation by cheaters, as exemplified by the well-known prisoner’s dilemma framework (also called *iterated prisoner’s dilemma* or IPD; 1). An important aspect of achieving mutual cooperation in biological settings is that cooperation is rarely “all-or-nothing” (2): players can alter their level of investment, i.e., how long they are vigilant or how long they groom. If cooperative investment becomes variable, individuals can adjust their amount based on their partner’s previous investment. Various strategies that utilize variable investment within IPD structured games have been proposed and, within a wide parameter space, cooperation is predicted to spread to fixation within the population (3–5).

One of the best known strategies proposed to yield stable cooperation in a continuous IPD (i.e. an IPD structured game with variable investment) is “raise-the-stakes” (RTS; 5). When applying the RTS strategy, an individual initially invests little with a novel partner to avoid being exploited (“testing the waters”) but gradually increases investment if the partner matches or exceeds it. In other words, players decide on levels of cooperation or investment based on a partner’s previous investment. This also implies that RTS does not increase if an investment is not matched by a partner, and cooperation only gradually increases over time when both partners cooperate (5). Under the model’s assumptions, the RTS strategy spreads quickly to fixation and cannot be invaded by non-cooperators once established (5). While the initial model does not specify cost and benefit functions, a follow-up by Killingback & Doebeli (3) shows a similar dynamic when the benefits of investment yield diminishing returns: the evolved cooperative ESS makes individuals start with levels that are below the payoff maximum, and responses to return benefits eventually lead to levels of helping that maximize benefit-to-cost ratios, which allows for no variation within a population. Both these models propose purely evolutionary strategies, i.e., the initial level of investment and updating rules are genetically coded. Furthermore, it is assumed that all individuals are in equal condition, that there is neither partner choice nor partner switching, and hence no scope for social bonding (also called “friendships”; 6) and payoff interdependence. At evolutionary equilibrium, all players therefore play the same strategy. These assumptions and outcomes contrast with a recent model by Leimar & Bshary (4), which also assumes that individuals can make variable investments to help someone, but does account for individual variability of players, with players learning about other individuals (where learning parameters may evolve according to selection on resulting behaviour). Furthermore, the model incorporates the main assumption of biological market theory (8) that individuals may choose among potential partners. Finally, the model allows for the development of social bonds, which in turn affects decisions, thus adding a key aspect of stable social life that biological market theory is silent on. The model predicts that, when encountering new, unrelated group members, players follow a strategy that may be called “all-in” (labelled ‘making friends’ by Trivers; 9): individuals start with high initial investment (4) – opposite to the predictions of the RTS. Individuals can subsequently change their investment based on where they stand with their partner: investment remains high if a partner asking for help has helped more than received, but investment drops if the partner is in debt. Debts may build up because of chance events but also because lower-quality individuals are more likely to need help.

Testing which decision rules wild animals use when starting a new relationship, and how they adjust levels of reciprocal investments based on the interaction history, is challenging, as it requires observing each pairwise interaction from the very beginning. We are not aware of any existing empirical study on wild-living populations that contains such data. While translocated cleaner fish, *Labroides dimidiatus*, show initially high investment to build up new relationships with ‘client’ reef fish (10), the study system does not fit a key assumption of the models, since the strategic options of cleaners and clients are asymmetric (only cleaners need to decide about levels of investment). Carter et al. (11) provide an example on captive vampire bats that to some extent fits the RTS dynamics: individuals go through a phase of exchanging low-cost grooming before they start with higher-cost food sharing. However, their framework treated food donations as binary events rather than variable amounts as assumed in the model. Other, more recent studies on captive vampire bats (12,13), as well as on zebra finches (14), underscore the diversity of currencies and temporal scales in forming cooperative bonds, highlighting the need for multivariate, longitudinal frameworks in free-ranging populations.

Allogrooming has long captured the attention of primatologists (15,16). Grooming is a key social currency and is often used as a proxy for pair bond strength (17). Studies using grooming interactions among primates to test theoretical cooperation models highlight that exchange dynamics may differ between short-term/immediate reciprocation (18–20) and intermediate-term and long-term exchanges. For example, in both macaques and capuchin monkeys, short-term reciprocity was observed but longer-term grooming preferences persisted independently of immediate exchanges (18,20). A study on wild-living baboons attempting to apply the logic of the RTS model found no evidence of adult females adjusting grooming efforts to what they had received from the female groomee during their last exchange (21). However, focal individuals had known each other long before data collection started (females are the philopatric sex in baboons) and were genetically related to varying degrees, therefore not fitting key assumptions of the model (12). Grooming exchanges in primates thus potentially offer a valuable context for studying cooperative strategies. However, a complication for the interpretation of grooming exchanges is that grooming can be traded not only for itself but also for other commodities (19,22–28). Models show that subordinates offering grooming in exchange for tolerance given by dominants may indeed yield stable cooperation (27,29). In the present study, we studied grooming to evaluate how exchanges between newly-met dyads of unrelated individuals develop over time. Specifically, we examined how newly immigrated individuals build grooming relationships with other group members.

Vervet monkeys (*Chlorocebus pygerythrus*) provide a suitable system to quantify how grooming exchanges develop from scratch over time, in order to test variable investment strategies such as RTS and “all-in” (4,5). Vervets live in promiscuous multimale/multifemale groups with regular male dispersal (30) and philopatric females forming the social core of the group (31–33). The females live in strict matriarchal hierarchies (32), where the rank between females and their offspring is determined by kinship: the last-born sibling gains the rank position right below their mother (34). Males, however, leave their natal group once they reach maturity (around 4-5 years) and continue to disperse every ~2 years (30,31,35). Males generally gain their rank through male-male conflict (32), but studies suggest relationships with resident females influences male rank as well (36), possibly at least partly due to females’ ability to resist mating attempts (36,37). Female vervets are known to trade grooming for feeding tolerance and coalitionary support (26,38), as well as potentially access to resources (25), with exchange values modulated by bond strength and correlated kinship. To exclude any effects of relatedness, our study on the development of grooming exchange patterns focuses on adult male-adult female (hereafter: male and female) dyads following male immigration into a new group. We only consider male-female grooming exchanges in this study as male-male grooming exchanges are rarely observed, especially between newly immigrated and established males. Female-female grooming exchanges are not of interest here, since females living in the same social group are often related and have prior social relationships – or at least knowledge of each other – thereby not fitting the key assumptions of both the RTS and “all-in” strategy. Having access to multiple groups habituated to human observers, we could observe males migrating between study groups. Because researchers are with groups six days a week, new but habituated males have been observed immigrating into groups and can thus be observed from their first days in their new groups. This allowed us to observe their grooming exchanges with novel, unrelated partners.

Under the assumption of long-term reciprocity and grooming being traded for grooming, we expect long-term grooming investment to be equal between partners. If both parties follow the RTS strategy, we expect low initial investment, due to the uncertainty of their partner (the “testing the waters” principle, 5). However, if they follow the “all-in” strategy (4), we expect high initial investment from both partners (Figure 1). For both strategies, the existing models predict that grooming services should be closely matched if parties respond to their partner’s investment. However, in our empirical case, there may be an important additional dimension, as interaction partners belong to opposite sexes. In biological market terms, it seems likely that grooming is not exchanged within one class of traders (as the models assume) but between two classes of traders with different life histories, which may also exchange other services. For example, grooming exchange patterns may be affected by females already having an established social network (26,31–33) while males may benefit from inclusion in the group (39) or are generally looking to increase their centrality, which positively affects survival and reproductive opportunities (40–42). Furthermore, females are generally the choosy sex in reproductive interactions, potentially causing males to provide more grooming in exchange, especially during the mating season. Contrarily, male primates are known to provide a variety of services to their group, including vigilance, predator mobbing and fighting neighbours (43–47). Importantly, although these asymmetries may cause systematic asymmetries between the sexes in grooming provisioning, they should not affect the models’ predictions regarding trends in grooming investments over time.

**Figure 1.**
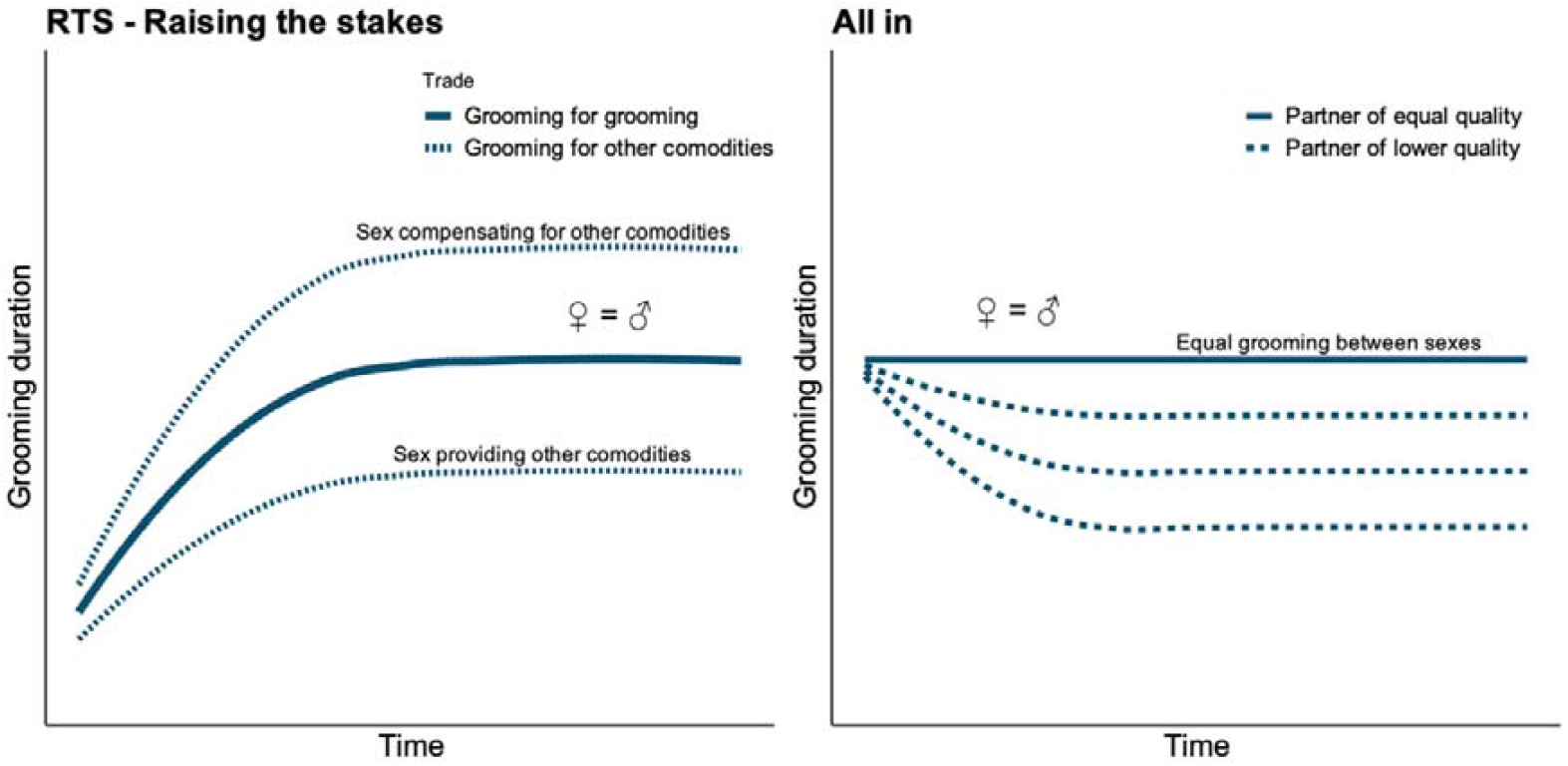
Predicted grooming investments based on the “raising-the-stakes” (RTS) and “all-in” strategies. In both predictions, males and females invest equally. When players apply “raising-the-stakes”, we expect grooming duration to start low and increase over time, until plateauing. However, if one sex is trading grooming for another commodity provided by the other sex – and assuming these commodities stay levelled – grooming duration might plateau at different levels. When players apply the “all-in” strategy, we expect a high initial grooming investment and a decrease in grooming duration when one of the sexes fails to reciprocate, due to being of lower quality.

Due to the logistical constraints inherent to field studies, our analyses on how grooming exchanges develop between females and newly immigrated male vervet monkeys focused on global grooming patterns rather than tracking precise dyadic exchange trajectories from their beginning. Thus, our results emphasise overall trends in male-female grooming exchanges over time rather than detailed pairwise grooming exchanges. We assessed whether reciprocity between newly immigrated males and females develops and compared this with the grooming exchanges between females and males with at least a year of tenure, as well as within males for which we have data following immigration and a year later. Specifically, we tested whether these grooming exchanges and their progression aligned with the predictions of the RTS (low initial investment, 5) or the “all-in” strategy (high initial investment, 4). By examining broader patterns of initial investment and reciprocity dynamics, this study provides insights into general cooperative relationship formation, highlighting the diversity of parameters that we need to consider for a proper understanding.

## Methods

We used observational data collected at the iNkawu Vervet Project (IVP) field site, Mawana Game Reserve (MGR), South Africa. MGR is a private game reserve of approximately 11000 ha, located in northern KwaZulu-Natal province. IVP data are collected year-round by trained field assistants who undergo monkey’s identification and interobserver reliability tests with the onsite scientific manager before collecting data. Over a period of two years (June 2022-June 2024), observers conducted three 20-minute focal observations on each adult individual in 10-day periods, though never twice on the same day or during the same time block (morning, noon and afternoon). Within focal observations, a grooming interaction was recorded for as long as the same two partners performed interactive behaviours with each other (excluding the duration of “behavioural gaps”, such as a bout of self-scratching, lasting less than 60 seconds), when both partners did not interact with a third party in the meantime and when neither partner moved significantly between behaviours. The durations of these interactions were considered in seconds for assessing the duration females groom males and vice versa. We used data from seven neighbouring groups of vervet monkeys. Because not all groups have been followed since the start of IVP in 2010 and habituation levels differ between the groups, group identity was considered in our analysis.

Because male vervet monkeys disperse multiple times during their adult life (30), we had numerous opportunities to assess grooming exchanges between novel partners. Based on long-term records – from the beginning of the project in 2010 – we excluded males that dispersed into the same group twice. Second dispersal into the same group was rare and always occurred within a year of a male’s departure. Because of these long-term records, we can say with high certainty that the males that were used in our study as novel partners had not been in their relevant study group before and therefore had no prior knowledge of individuals within those groups. Because some males dispersed into our population from unknown groups, and were therefore unhabituated, each male’s “habituation level” (i.e., habituated vs. unhabituated) was considered. Individuals were considered habituated when they could be observed from a 5-10-meter distance by researchers without altering their behaviour, showing signs of distress such as self-directed scratching or looking at the researchers. If a male was born in our study population or had spent time in a habituated group previously, he was considered habituated (as these individuals generally are unbothered with the presence of humans). If he was not, he was marked as unhabituated and excluded from the dataset, to ensure accurate data collection on grooming duration. A total of 15 males were thus excluded from the dataset, constituting 202 grooming observations with females. Males that had been in a group for over a year were considered habituated.

Our first step was to consider grooming reciprocity, since both tested models predict stable reciprocity over time (both strategies adjust investment based on their partner’s investment). To account for grooming performed by both sexes, we assessed reciprocity during the first year after a male’s immigration – as we aimed to study exchanges among novel partners. Reciprocity was calculated as a continuous variable ranging from −1 to +1, where 0 represents equal grooming effort by both partners, using the following formula:

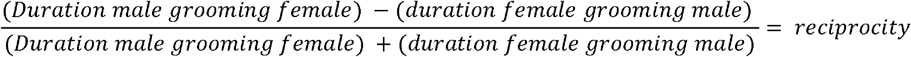

A positive score reflects greater male grooming effort, a negative score greater female grooming effort. The duration of grooming interactions were summed per male-female dyad, per observation day. Since there was a maximum of 20 minutes observation time per male, all grooming interactions with female partners in those 20 minutes were directionally summed. It was uncommon for a male to have more than one female grooming partner per 20-minute observation (39 out of 220 focal observations).

This calculation, however, does not discriminate between cases where grooming, while unidirectional, differs greatly in duration (e.g., a male grooming a female for 10 seconds and for 100 seconds both get a score of +1). As a result, although the index is formally based on grooming duration, because grooming interactions in vervets are often one sided (~72% of recorded grooming interactions), this index effectively captures both the duration and frequency of grooming bouts. To verify this, we also calculated a frequency-based reciprocity index, based on the number of grooming bouts performed by each partner per day of male residency. The two metrics were highly correlated (*Pearson’s r* = 0.94, *p* < 0.0001), indicating that daily grooming effort and frequency varied similarly as males’ tenure increased. We therefore proceeded with the duration-based index. While we acknowledge that grooming efforts with different durations are not equal, our analyses focus on the potential change (slope) in grooming exchange over time, and this index captures that pattern adequately. To simplify the modelling process, we transformed the data into a binomial format. Grooming exchanges between dyads over a day that had a positive reciprocity score (male-biased effort) were considered as 1, and exchanges with a negative score were considered as 0 (female-biased effort). Although we acknowledge these choices may limit nuance, these decisions were necessary simplifications given data limitations and distribution challenges. In total, 26 out of 78 newly immigrated male-female exchanges did not have a value precisely of −1 or 1 and were rounded up or down. The graphic projections of the data revealed that both binomial and continuous but bound reciprocity measures exhibited similar patterns.

Our reciprocity curve showed clear unequal reciprocity in the first months of the new relationship (see Results, Figure 2). We therefore continued our analyses wanting to uncover which partner adjusts their behaviour until they reach stable reciprocity. To do this, we first had to make a distinction between novel partners (newly immigrated males) and more established partners. We therefore aimed to create two categories of males with an equal length in days. We chose to compare newly immigrated males with males that had already been present in a group for at least a year, to make sure they were sufficiently familiar with their female partners. To find the period in which there was unequal investment and the point in which this inequality changed we used a breakpoint model, a statistical method used to identify points of abrupt change (48,49), implemented in the “segmented” package (50). This breakpoint model included day of group membership of the male, season, and group as covariates. This gave us a specific day in a male’s tenure in which grooming reciprocity changed, which we used to define our male categories.

**Figure 2.**
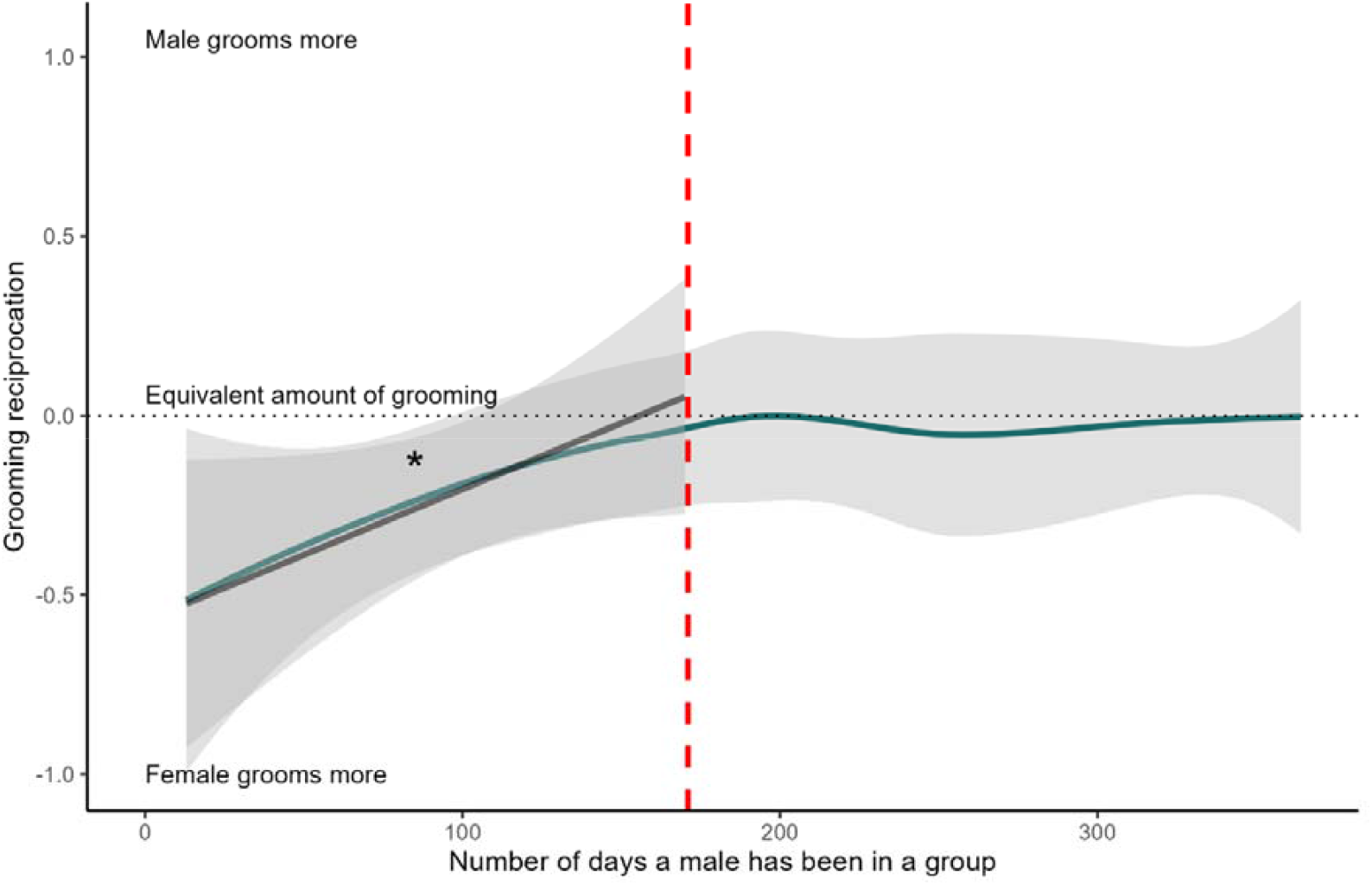
Grooming reciprocation rates between males and females over the first year of a male’s tenure. Positive values indicate males investing more and negative values females investing more. The red dotted line shows the breakpoint found by the breakpoint model. Therefore, any exchange with a male to the left of this line is qualified as an exchange with an immigrant. Reciprocity increases significantly for these exchanges (*indicated with*).

After defining the breakpoint in our model, we compared initial investments of females towards novel partners (immigrant males) and vice versa, by comparing the duration of grooming bouts as well as the frequency of exchanges between both partners. We compared these initial investments with later investment, i.e., investment between females and established partners (males with >1 year of tenure, hereafter called residents) and the progression of investment over time. The use of this breakpoint to delimit the period in which a male is still novel to female partners should not be considered as a general benchmark for vervet immigration but instead be treated as a data-derived marker of behavioural change in our dataset. We acknowledge that social integration is unlikely to follow a discrete or universal temporal threshold but instead represents a gradual process that can be determined using different behavioural metrics, which will inevitably identify different breakpoints. Literature on male immigration period in vervets is scarce, and although a 60-day cutoff was used twice previously (51,52), this thresholds appear to be largely conventional and is not directly empirically grounded. Applying such a cutoff to our data would not alter our conclusions but likely strengthen our observed contrasts – at the expense of statistical power. Importantly, identifying a breakpoint in the development of male-female grooming exchanges is itself a key result, yielding as a by-product a biological category of “immigrant” or “resident” male based on social behaviour. We use these terms to follow the general vocabulary in the field, while the labels “first-year vs second-year males” or “novel vs established partners” are viable alternatives.

All data were analysed using R (v4.2.2), using generalized linear mixed-effects models within the *glmmTMB* package (53), allowing us to include complex random effects structures. The binomial response variable, *reciprocity*, was modelled as a function of the status of the male (either immigrant or resident) in an interaction with the number of days the male has been in his status (ranging from day 0 to the day found by the breakpoint model for both male categories) as a predictor variable. Since both the mating season as well as potentially lactating females are known to influence male-female relationships (54,55), we divided the year into four seasons of equal length: baby season (the three months when most births occur), mating season (three months when most mounts occur) and two three-month periods in between (defined here as winter and summer, distinguished by their contrasting food availability), and included season as a fixed effect. Data was insufficient to include the male-female dyad over the observation day as a random effect structure, which caused convergence problems for our model. Instead, we included a nested random effect structure of male and female identity to account for repeated observations within individual males, as well as dyad-specific variation in grooming duration between specific male-female pairs, and the group identity. A table of all models can be found in the supplementary materials S1. Significance of fixed effects was evaluated using Type II Wald χ^2^ tests, and effect sizes were interpreted using model coefficients and emmeans-based post-hoc contrasts.

To analyse grooming durations, which were continuous, strictly positive, and right-skewed, we used a Gamma distribution with a log-link. Residual diagnostics (via DHARMa) supported the suitability of the model. For the analysis of grooming frequencies, the same structure was used but performed with a binomial distribution.

Directional grooming bout duration (in seconds) between a male and female was our response variable to test grooming duration. When two individuals groomed each other twice on a day but separated by more than one minute, an interaction with a third individual or travel, the interactions were considered as separate bouts. We analysed female-given and male-given grooming in two separate models. For each sex-specific dataset, grooming duration was modelled as a function of the interaction between the male’s status (immigrant vs. resident) and the number of days the male had been in that status (for both male categories ranging from 0 to the breakpoint), with season included as a fixed effect. The day in the status was z-score transformed and the two-level factor status was coded using sum-to-zero (effect) contrasts, to enhance interpretability of the results of the interaction term (56). Manual effect coding of the status was not feasible because dyads and male-female combinations – present in our random effect structure – did not occur in both status levels, producing non-overlapping levels within random-effect strata, thus preventing model estimation. We used a random-effect structure tailored to each sex-specific dataset: both models included a nested random effect structure of male and female identity to account for repeated observations within individual males, as well as dyad-specific variation in grooming duration between male-female pairs. In addition, to allow groups to vary temporally in grooming investment, we included a random slope of the day of a male’s tenure within group. Model convergence for this random-slope structure was achieved by using the BFGS optimizer. Since ten males in the dataset were present as both immigrant and resident, we performed a within-individual comparison as well, by performing the same model with the subset of these ten males. Results of this can be found in the Supplementary materials, Table S3c,d and are consistent with the full data model.

For the grooming frequency dataset, we noted whether there was a male-female or female-male grooming interaction (yes/no) observed within any focal observation performed on one of our study males, creating two outputs per focal observation (male-given grooming, female-given grooming). Although this method did not allow for the inclusion of multiple grooming events in the same direction to be recorded, because of our definition of a grooming interaction (no interaction with a third party, significant movement or more than a minute break in between grooming behaviours) these observations were rather scarce (22% of cases where a male was groomed more than once by the same or different female partners; 22% of cases as well where a male groomed one or multiple female partners more than once). Our binomial models had the same structure as our grooming duration models, with the frequency of grooming exchanges as the response variable and an interaction between the status of a male (sum-coded) and the number of days a male has been in his status (z-transformed), analysed per grooming sex, season as a fixed effect, and including the male over his tenure day and group identity as random factors. Because of our binomial structure, female identity could not be included since they are not applicable for focal observations where there was no interaction observed with a female.

The rank of newly immigrated males could not be reliably calculated due to the lack of conflict data recorded in the short period they were with their new group. Similar problems occurred for the inclusion of a male’s age, since many males were not born in our population and therefore impossible to age reliably. While newly immigrated males might have had mating opportunities, their potential offspring generally has not been born yet, as the average gestation period of female vervets is around 5.4 months (57). Males with over a year of tenure, on the other hand, have had at least one mating season and could therefore have fathered offspring. Due to the promiscuous nature of vervets (e.g., 54), they are considered potential fathers. Thus, if included in our models the potential of being a father would show an almost perfect overlap with tenure length. We therefore decided not to include any individual-specific parameters, but rather approach our question from a more global, group perspective – besides accounting for the male’s identity in our models. However, a brief analysis was performed on the subset of males of which we were able to calculate rank scores and their grooming durations with females. Note that the possibility to calculate rank scores is dependent on several factors (how long the male has been in a group, how central he is, how many conflicts can or have been recorded, etc.), which might bring different biases into the subset. The results of this analysis – consistent with our main results – can be found in the supplementary materials (Table S5).

## Results

To determine whether grooming exchanges between immigrant males and adult females followed the “raise-the-stakes” (RTS) or “all-in” strategy or other dynamics, we first looked how the grooming reciprocity index developed between newly immigrated males and females. Data were based on 18 males and 73 females. Initial investments in exchanges with novel males were highly female-biased, while grooming reciprocity became more balanced over the first year but remained slightly biased towards females investing more (Figure 2; reciprocity over one-year period, with negative values meaning that females groom more and positive values meaning that males groom more: *mean* = −0.10, *median* = −0.23). The segmented regression model revealed a significant breakpoint at day 171 (± 35.2 SE), indicating a change in grooming reciprocity over time. The effect of day before the breakpoint was significantly positive (β = 0.015, SE = 0.006, z = 2.45, p = 0.014), suggesting an increase in reciprocity with male tenure. After the breakpoint, the slope changed direction and became negative (U1.Day: β = –0.020, SE = 0.009, z = –2.26), indicating that reciprocity decreased after the breakpoint. This confirms a change in the trajectory of grooming relationships, consistent with our initial visual inspection of the raw data. The results hence show that males and females do not use the same strategies during the development of grooming exchanges, warranting a more detailed analysis of what each sex is doing.

For the relevant analyses, we decided to investigate i) what happened until the breakpoint, and ii) compared the dynamics with those of females exchanging grooming with males that had already been in the group for one year, to control for seasonal effects on behaviour. For simplicity, we call males until the breakpoint ‘immigrants’, and the males that had been in for a year ‘residents’, though we acknowledge that these terms should not be seen as categories. As mentioned before, we used data on 18 immigrant males, of which 10 males were present in both the immigrant and resident period (Table 1). Over our two-year data collection period, consisting of 272 observation hours on both immigrant and resident males, we recorded 405 grooming events involving females and immigrant or resident males (respectively 112 and 293 events). Immigrant males groomed females in 61 recorded instances (mean duration: 93.06 seconds), whereas females groomed immigrants 51 times (mean duration: 158.60 seconds). Similarly, residents groomed females 142 times (mean duration: 137.42 seconds), while females groomed residents 151 times (mean duration: 157.70 seconds). When comparing the slopes of reciprocity of immigrants and residents, we found a significant positive slope for immigrants (*estimate* = 0.87, *SE* = 0.35, *p* = 0.01), but no significant slope for residents (*estimate* = 0.33, *SE* = 0.26, *p* = 0.20). However, both slopes did not significantly differ from each other, since both were moving in the same direction (*estimate* = 0.55, *SE* = 0.37, *p* = 0.14). Resident males furthermore had a higher reciprocity than immigrant males (χ^2^ = 6.05, *p* = 0.01; Table S2). The random effects explained approximately 15.4% of the total variance on the latent (logit) scale, with almost all attributable to the male identity (Table S2).

**Table 1:** overview of each male category and from which day they are in a certain category.

| Male category | Day of residence | n males | n female partners |
| --- | --- | --- | --- |
| *Immigrant* | 0 – 171 | 18 | 39 |
| *Resident* | 365 – 536 | 22 | 61 |

The interaction between male status and days in status significantly explained variation in female-given grooming duration (χ^2^ = 7.46, *p* = 0.006; Figure 3, Table S3a). Simple-slopes analysis showed that females decreased grooming of immigrant males over time (*estimate* = −0.35, *SE* = 0.12, *p* = 0.004; Figure 3, Table S3a), whereas grooming of resident males remained stable (*estimate* = 0.06, *SE* = 0.11, *p* = 0.55). The main effect of status and season were not significant (Table S3a). Random effect estimates indicated that dyad identity accounted for most variance in grooming durations, whereas group-level variation and random slopes over a male’s tenure were near zero.

**Figure 3.**
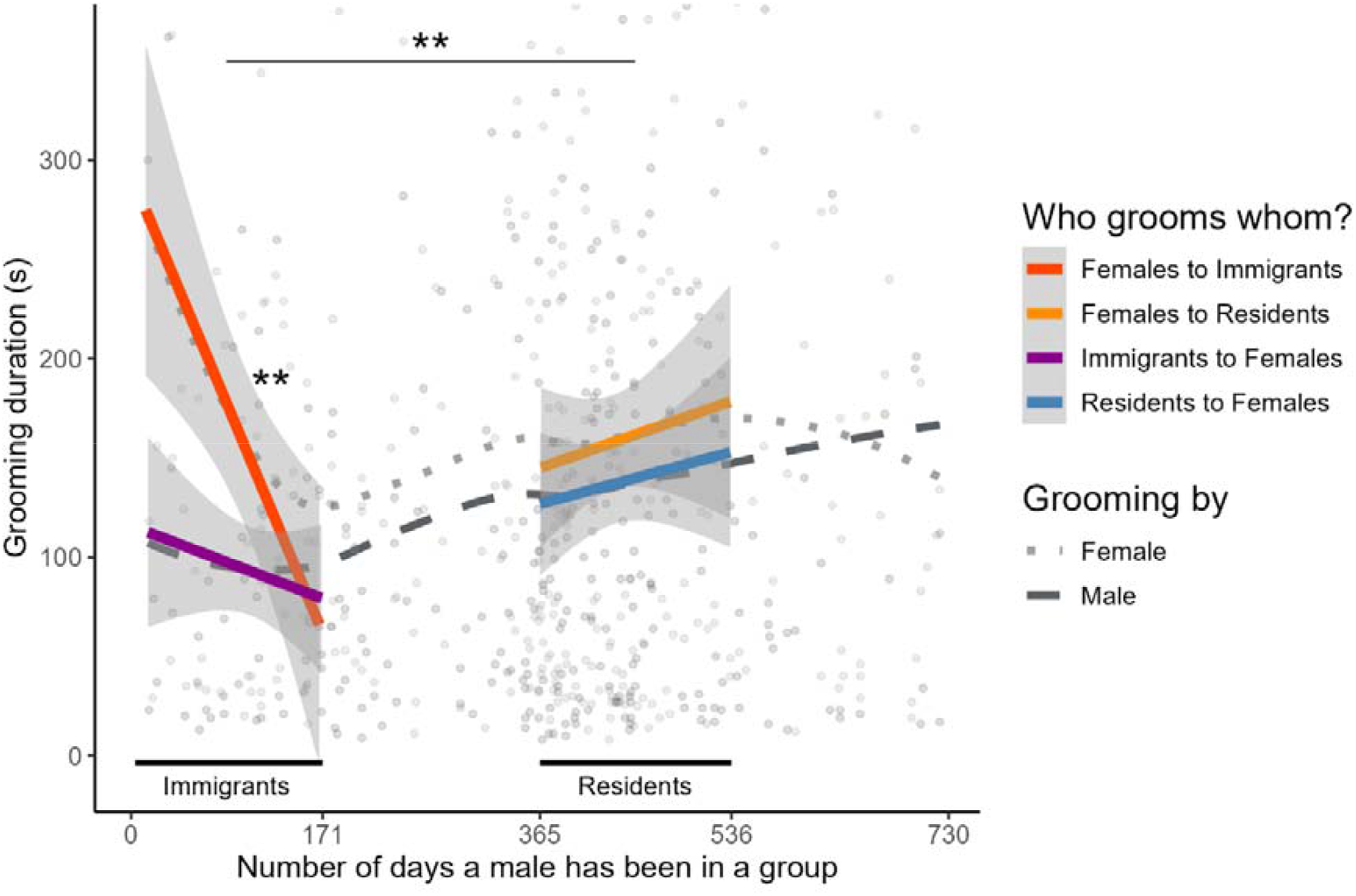
Grooming duration between females and males in relation to the male’s status (raw data). The dotted gray lines represent the overall trends for grooming duration, with the smaller dots for females and larger dots for males. Raw datapoints of female-given grooming are in light grey, male-given grooming are indicated in dark grey. The colored lines show model trends for different grooming exchanges: females grooming immigrant males (dark orange), females grooming resident males (yellow), immigrant males grooming females (purple), and resident males grooming females (blue). The decrease in grooming duration by females grooming immigrant males is significant (*indicated with* ^**^). Females also changed their grooming duration over time differently for immigrants than they do for residents (*indicated with* ^**^). All other slopes were not significant.

Male status showed a marginal effect on male-given grooming duration (χ^2^ = 3.84, p = 0.05). Although the model estimated longer grooming by resident males, the pairwise contrast was not statistically significant (*estimate* = −0.11, *SE* = 0.08, *p* = 0.195). All temporal effects, including the interaction with status, were not significant (Figure 3; Table S3b). Season had no effect, and random-effect variance was negligible at the group and dyad levels, with modest variation attributable to male identity.

For female grooming frequency, there was no evidence that male status, time in status, their interaction or season affected grooming rates (Figure 4; Table S4a). Random-effects variances were minimal, indicating little individual or group-level structure. Male grooming frequency, however, showed a significant main effect of male status: immigrant males groomed females less often than resident males at the average day (χ^2^ = 7.65, *p* = 0.006; Figure 4, Table S4b). No temporal effects or interactions were detected, season had no effect and random-effect variance was again negligible.

**Figure 4.**
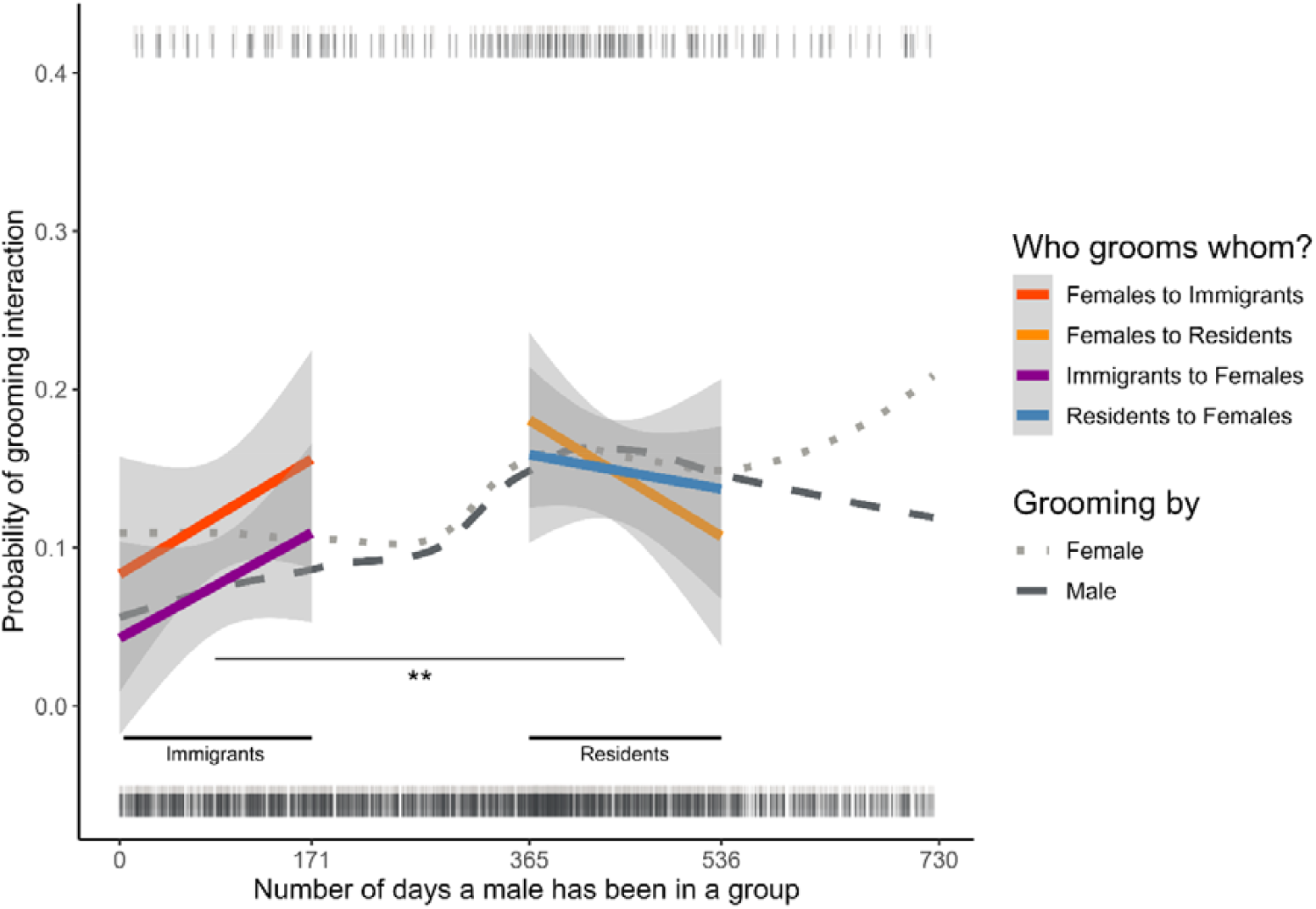
Grooming frequency between females and males in relation to the male’s status (raw data). The dotted gray lines represent the overall trends for grooming frequency, with the smaller dots for females and larger dots for males. The colored lines show model trends for different grooming exchanges: females grooming immigrant males (dark orange), females grooming resident males (yellow), immigrant males grooming females (purple), and resident males grooming females (blue). The difference in grooming frequency by immigrant and resident males grooming females is significant (*indicated with* ^*^). All other slopes were not significant. Raw datapoints are indicated by the dark (male grooming) and light (female grooming) grey ticks at the bottom (value of 0, no grooming present) and top (value of 1, grooming present) of the plot.

## Discussion

To assess how males and females build personal relationships in a monkey species with high levels of allogrooming, we examined how grooming exchange patterns between immigrating males and philopatric female vervet monkeys develop over time. While the two evolutionary models we tested both predict stable reciprocation of grooming investment, we found a strong female-bias in grooming investment in the first months of a new relationship. We found that females and males showed rather opposite patterns in their grooming investments. Males provided practically constant grooming services to females in terms of grooming duration (fitting the “all-in” prediction) but increased their grooming frequency after a year of residence (fitting the RTS prediction). In contrast, females showed the highest grooming investment in terms of duration with newly immigrated males, while grooming them as frequently as resident males; their investment in duration decreased over months until stabilizing at a level that was similar to male grooming investment into females. While the female data are thus more in line with the “all-in” strategy (4) than with the RTS strategy (5), the asymmetry suggests that females and males follow different strategies. We therefore need to explain why females initially groom newly immigrated males for longer and only lower their investment over several months to eventually closely match male investment. We discuss these results in more detail below.

According to the RTS strategy, initial investment of both parties should be low, and investment increases when the partner reciprocates (5). Neither sex showed any evidence that initial grooming bouts given are short and then slowly increase over time if matched. Instead, grooming bouts either started with a particularly high investment in duration (male immigrants grooming females mean duration: 93.06 seconds, females grooming immigrants mean duration: 158.60 seconds) in the beginning with a decrease over time (females) or were rather stable over time (males). While females stayed stable in their grooming frequency towards males, resident males seemed to groom females more frequently than immigrant males. This latter result matches the RTS hypothesis, but only if we accept that a step increase in grooming frequency after a year may be still called “raise-the-stakes”. The aspect of RTS that grooming exchanges build up in a reciprocal way over several steps can be refuted despite an important shortcoming in our data: we do not have enough observations to analyse the precise dynamics of the progression of grooming exchanges in specific male-female dyads. Although we corrected for the repeated observations of grooming individuals in our models, we could not test whether receiving a long or short grooming bout is followed by a matching return grooming bout, as would be expected if individuals use a decision-rule that is based on “give as much as you get”. Nevertheless, the data suggest that for male-female vervet monkey grooming exchanges, females are not concerned about whether male partners return their grooming investments in the form of grooming. In other words, the risk of being “cheated” does not appear to affect female grooming decisions, characterised by their extra high grooming investment. Thus, it appears that females groom new males for other reasons. Females and established males groom each other for rather similar durations, suggesting that once males are fully resident, grooming is largely traded for grooming.

The model by Leimar & Bshary (4) predicts that the first act of helping reflects an investment as close to the maximum an individual is willing to give, and that such help is readily given to a new partner as long as this individual is or becomes a group member. The female results partly fit the model’s predictions in that initial grooming investments in duration are as high or even higher than later grooming investments. As discussed, we lack the detailed dyad-level data necessary to test how individuals respond to a build-up of grooming asymmetries within dyads, to assess whether decision-rules follow the logic of reciprocity, but the high levels of variance explained by the dyad as a random effect suggests little changes of grooming over time between partners. However, we consider it unlikely that the model can explain why there is an initial grooming bias by females that eventually disappears, and close-to-matching grooming exchanges develop. More precisely, we consider it unlikely that females reduce their initial grooming investment simply because they do not receive adequate matching by the male partners. If this were the case, females would make systematic errors every time they encounter a new male. Such errors should be eradicated by learning or natural selection, leading females to start with lower initial investment when encountering new male partners. In fact, females give newly immigrated males especially long grooming services, longer than what they receive back from these partners. Therefore, other factors need to be considered to explain the initial asymmetry in male-female grooming exchanges.

The grooming bias cannot be explained by differences between the sexes regarding the need to form new social relationships. In vervet monkeys, adult females form the social core of the group, being surrounded by related females with whom they have established social bonds (32). Such female-female bonds allow females to often dominate males, despite being smaller (59,60). Thus, females do not need to form new strong relationships, whereas immigrant males do. One could therefore have expected that immigrant males show high initial grooming investment into females in order to better integrate into the group through the formation of social bonds with resident females (31–33). Immigrant males could also benefit from grooming females more by bringing increased mating opportunities (39,61,62) and a rise in the male hierarchy due to agonistic support by females (36,42,63). However, immigrant males groom females for as long as established males do (fitting the “all-in” hypothesis) but less frequently as established males do (fitting the RTS hypothesis). Clearly, immigrants do not show extra effort, whereas females do. If the females’ effort were limited to the first few days, one could have hypothesized that a likely higher ectoparasitic load in newly immigrated males (64) might have caused females to prolong the grooming of immigrant males as a way to obtain food. However, it seems unlikely that immigrants’ parasite loads take months to decline.

Two factors may explain the initial asymmetries in grooming. First, females seem to expect long-term benefits from grooming immigrant males, as they start with a higher initial investment and do not get paid back directly in the form of grooming. There are several hypotheses as to why having new males join a group could yield net benefits for females, despite increased feeding competition (65). Among them are inbreeding avoidance (66,67) and the possibility that having a more male-biased adult sex ratio in a vervet group may promote female dominance and hence increased partner choice (68), as is predicted by DomWorld (60). Complementary benefits may arise because additional males increase the total amount of so called ‘*male services*’ (69). As it stands, primate males may help in territorial defence (22,43,45), yielding increased access to resources (70) and decrease predation risk due to vigilance and active defence (69,71). Second, being the dispersing sex, male vervet monkeys can in principle choose among several groups, which would cause competition between different female core groups provided that additional males provide net benefits. This asymmetry in group-level partner choice options in a biological market (8) thus contrasts with the individual level, where vervet females choose among males (37). In our scenario, once males are convinced to stay, females can reduce their grooming investment to “normal” levels, perhaps because males are then potential fathers, likely to have an incentive to stay to protect their infants. A question arising from this scenario is how females share the initial investment of enticing males to stay among themselves. A “tragedy of the commons” scenario in which free-riding is the best option appears to be highly unlikely as even humans struggle to solve such situations in a cooperative way (72,73). Potentially, the game resembles more a volunteer’s dilemma (74) rather than an n-player prisoner’s dilemma, leading to contributions being under negative frequency dependence. However, contributions may also become self-serving as genetic relatedness among females may yield indirect fitness benefits of contributions through kin selection (75,76), and individualised direct benefits may be accrued if some benefits of early grooming are partly partner-specific (e.g., preferential tolerance or proximity to a particular male). A proper evaluation of these alternative scenarios will require additional data beyond the scope of this study.

We recognize several limitations. First, rapid initial escalation of grooming – if it occurs within hours of contact – may have eluded our sampling resolution, potentially conflating early “raise-the-stakes” dynamics with longer-term declines. However, our results remain relevant for longer-term dynamics because any short-lived initial escalation followed by a decrease would suggest a more complex strategy, not necessarily aligned with either RTS or “all-in”. Second, grooming is known to function as a ‘currency’ that can be traded for various other cooperative behaviours (e.g., food sharing, tolerance), as has previously been shown in this and other primate species (19,22–28). Because we did not measure these alternative currencies in this study, we cannot exclude the possibility that females invest in grooming immigrant males because these yield returns in forms we did not quantify. However, although grooming can be exchanged for other things, this cannot explain the decrease of grooming duration for females towards immigrants over the first six months. Thus, we consider it most parsimonious that the initial grooming investment by females functions to bind immigrant males to the group.

In conclusion, our results reveal a heterogenous pattern of how males and females groom each other over time. While a few aspects of these patterns fit the “all-in” strategy (4) or the “raise-the-stakes” strategy (5), the differences between female and male grooming patterns highlight the importance of considering initial asymmetries within cooperative exchanges when interpreting the data. Cooperation is rarely symmetrical in its payoffs in empirical settings whenever both partners might differ with respect to alternative choices (as formulated by biological market theory, 8), hold different social statuses, positions in hierarchy, embody differing life-history traits, or may offer different resources. The existing models were useful in yielding testable predictions on how new social relationships are built. New models that explore the importance of asymmetric variables will yield ever more realistic concepts.

## Acknowledgments

We thank the team at IVP for their crucial contributions to data collection, group monitoring and help with logistics, as well as Nokubonga Dhlamini, Michael Henshall, Siboniso Thela and Zonke Mbutho for running the field site as well as their invaluable support during data collection. We are grateful to the van der Walt family for permission to work in their reserve, as well as to Ezemvelo KZN Wildlife for granting us the ethical permissions to do so. We thank Radu Slobodeanu for his statistical assistance, as well as the anonymous referees and the associate editor, for their valuable suggestions on the manuscript.

## AI declaration

During preparation of this manuscript, the authors used AI to improve readability of the text and to assist in refining code. All AI-generated material was reviewed, edited, and verified by the authors. AI was not used for data interpretation or drawing conclusions. The authors take full responsibility for the accuracy and originality of all content.

## Author Contributions

R.B. conceived of the presented idea. E.vdW. contributed the data. J.A.T. and M.G.R. conceptualized data analysis, carried out the data cleaning and analytic calculations. C.vS. helped with the interpretation of the results. J.A.T. wrote the drafts of the manuscript that were edited by R.B., with further input of all authors. All authors gave final approval for publication.

## Data availability and open access

All relevant data and code are available on OSF: https://doi.org/10.17605/OSF.IO/AZYBR.

## Ethics statement

This research adhered to the Association for the Study of Animal Behaviour Guidelines for the Use of Animals in Research (Behaviour, 2018). The study was purely observational and involved no handling or experimental manipulation. Vervet monkeys are not a protected species in South Africa, and no permit or ethical approval number is required for observational research on this species conducted on private land. Nevertheless, the relevant local authority, Ezemvelo KZN Wildlife, and the van der Walt family, the owners of Mawana Game Reserve where the field site is located, approved the study and granted ethical permission. All individuals observed in this study were habituated to human presence and there were no interactions between humans and study subjects during the study.

## Electronic Supplementary Material

**Table S1:**
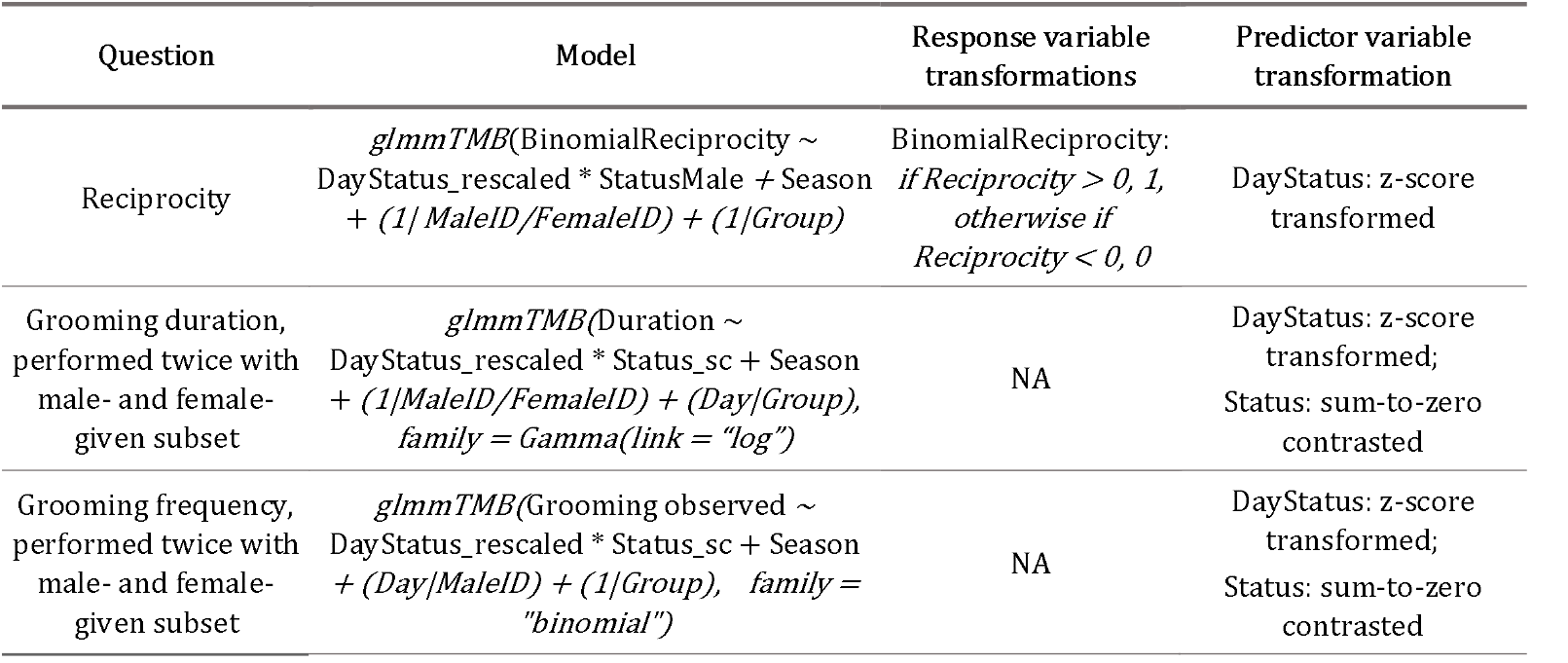
overview of statistical models applied to different datasets.

**Table S2:** Model output of the reciprocity model (binomial). The Est (95% CI) column reports odds ratios (OR) with 95% Wald CIs, obtained by exponentiating logit-scale estimates and their intervals. Fixed effects are shown as ORs; ratio repeats the OR (or the OR per +1 SD for slopes). Pairwise contrasts (Status; Season) are tested on the logit scale and presented as ORs with Tukey-adjusted CIs and p values; the z column is the Wald statistic returned by emmeans. Slopes (emtrends) for *DayStatus_rescale* are displayed as OR per +1 SD (exp of the logit slope); slope contrasts are differences in slopes on the logit scale (reported with z and p; ratio is not applicable). Random effects (SDs and correlations) are descriptive only. All tests are two-sided.

| Source | Term | Est (95% CI) | z | ratio | p.value |
| --- | --- | --- | --- | --- | --- |
| Fixed effects | (Intercept) | 0.200 (0.082, 0.485) | -3.560 | 0.200 | < 0.001 |
| Fixed effects | DayStatus_rescale | 2.394 (1.214, 4.721) | 2.520 | 2.394 | 0.012 |
| Fixed effects | StatusResident | 3.273 (1.444, 7.421) | 2.840 | 3.273 | 0.005 |
| Fixed effects | Season.L | 2.798 (1.170, 6.689) | 2.314 | 2.798 | 0.021 |
| Fixed effects | Season.Q | 0.605 (0.218, 1.684) | -0.962 | 0.605 | 0.336 |
| Fixed effects | Season.C | 0.882 (0.294, 2.645) | -0.225 | 0.882 | 0.822 |
| Fixed effects | DayStatus_rescale: StatusResident | 0.579 (0.279, 1.201) | -1.470 | 0.579 | 0.142 |
| Random effects | SD (Intercept) FemaleID:MaleID | < 0.001 |  |  |  |
| Random effects | SD (Intercept) MaleID | 0.774 |  |  |  |
| Random effects | SD (Intercept) Group | < 0.001 |  |  |  |
| Pairs (Status) | Immigrant / Resident | OR 0.305 (0.135, 0.693) | -2.840 | 1.357 | 0.005 |
| Slope (Status) | Immigrant | OR 2.394 (1.214, 4.721) | 2.520 | 2.394 | 0.012 |
| Slope (Status) | Resident | OR 1.385 (0.840, 2.285) | 1.277 | 1.385 | 0.202 |
| Contrast slope (Status) | Immigrant - Resident | 0.547 (-0.183, 1.277) | 1.469 | 0.579 | 0.142 |
| Pairs (Season) | Baby - Summer | OR 0.428 (0.043, 4.274) | -0.948 | 0.428 | 0.779 |
| Pairs (Season) | Baby - Mating | OR 0.228 (0.038, 1.382) | -2.108 | 0.228 | 0.151 |
| Pairs (Season) | Baby - Winter | OR 0.266 (0.067, 1.061) | -2.459 | 0.266 | 0.067 |
| Pairs (Season) | Summer - Mating | OR 0.533 (0.059, 4.816) | -0.734 | 0.533 | 0.883 |
| Pairs (Season) | Summer - Winter | OR 0.622 (0.076, 5.069) | -0.581 | 0.622 | 0.938 |
| Pairs (Season) | Mating - Winter | OR 1.167 (0.428, 3.183) | 0.396 | 1.167 | 0.979 |

**Table S3:**
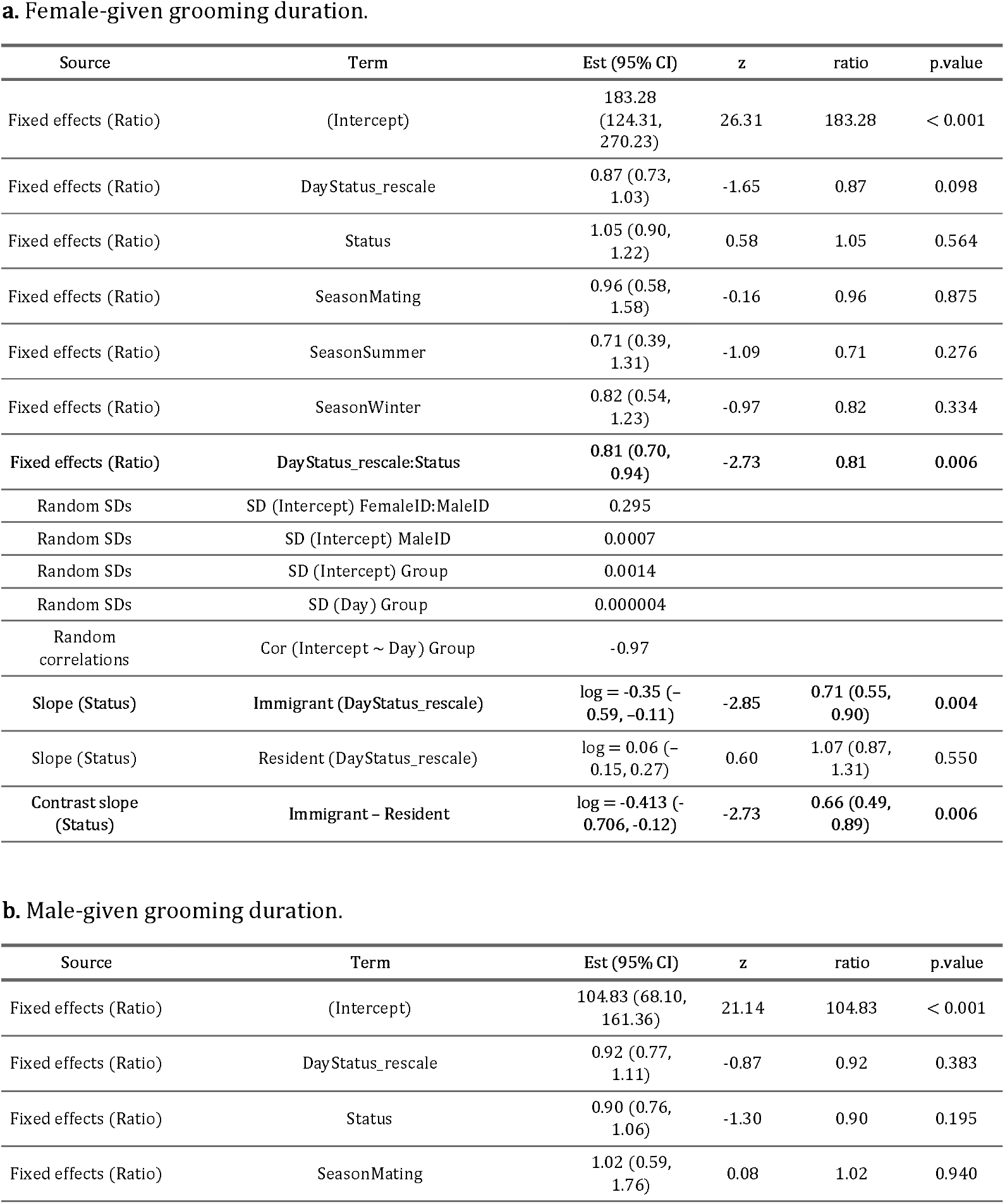
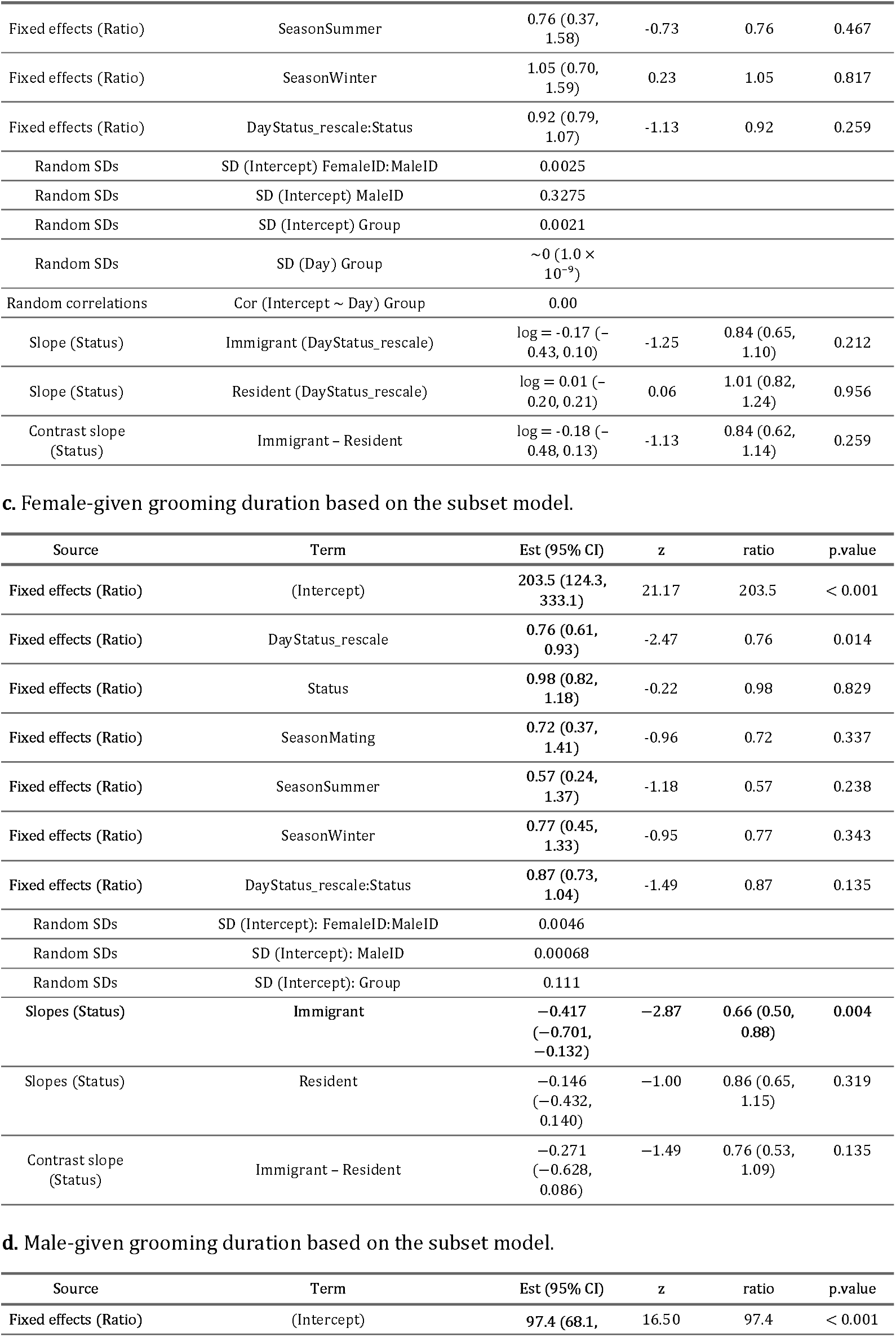
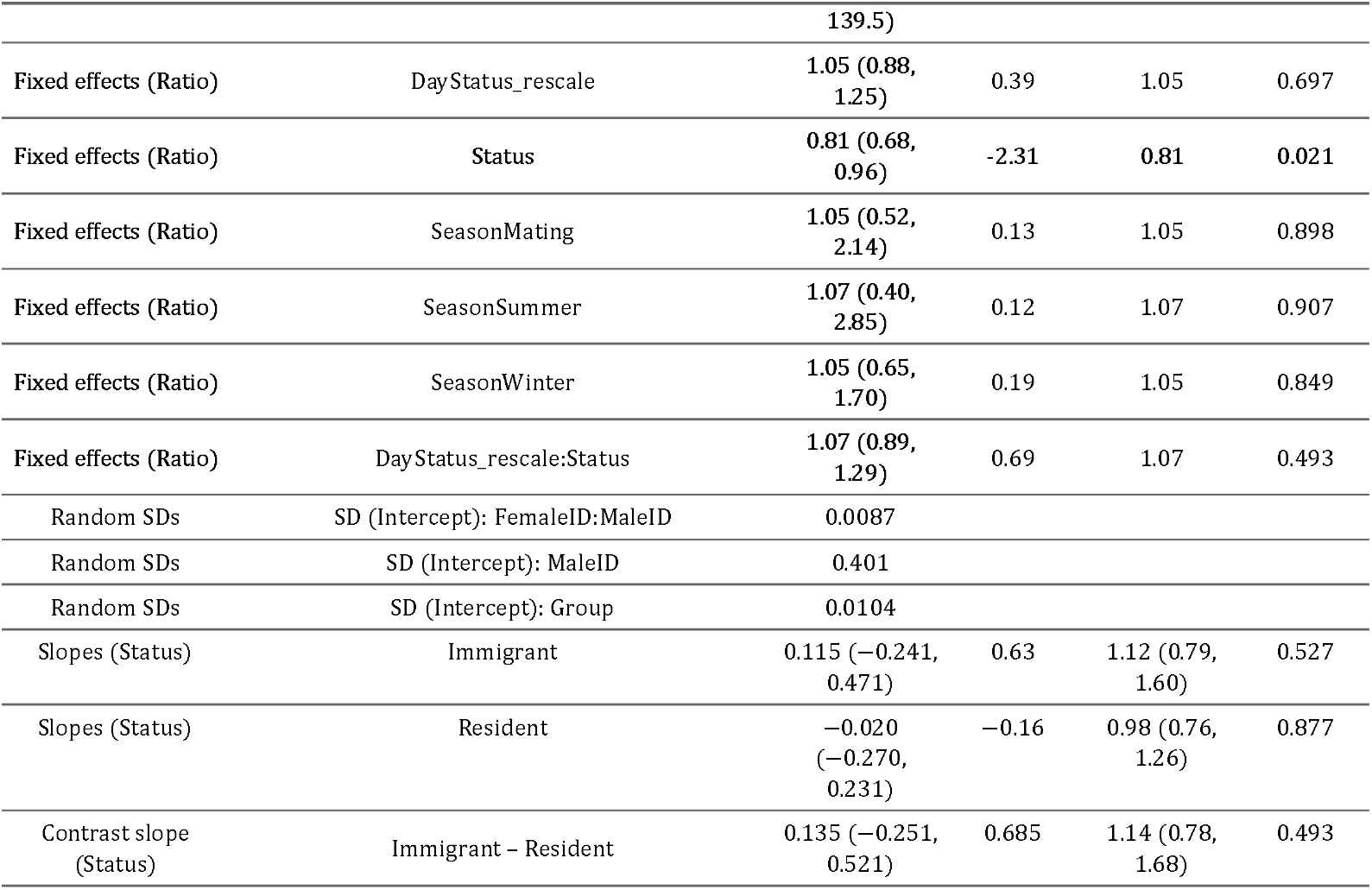
Model output of grooming duration models. Relevant results are in bold. Fixed effects are presented as ratios (exponentiated coefficients) with 95% confidence intervals, z-values, and associated *p*-values. Random effect standard deviations (SDs) and correlations (Cor) are reported on the model’s link scale. Slopes of *DayStatus_rescale* are given per +1 SD change, with slope contrasts reported on the link scale.

**Table S4:** Model output of the grooming frequency model (binomial). Relevant results are in bold. Entries list the effect name (“Source”) with point estimate and 95% confidence interval in the Est (95% CI) column. Fixed effects are reported as odds ratios (OR) (exponentiated logit coefficients). Pairwise contrasts (Groomer^*^Status; Season) are tested on the logit scale and presented as OR with Tukey-adjusted CIs and p-values. Slopes (Status|Groomer) for *DayStatus* are shown as OR per +1 SD of the standardized predictor; slope contrasts are differences in slopes on the logit scale (not exponentiated). The z column reports the Wald statistic returned by emmeans or the model summary; p.value is two-sided (Tukey-adjusted for multiple contrasts where applicable). Random effects (SDs and correlations) are descriptive only.

**a. Female-given grooming frequency.**
| Source | Term | Est (95% CI) | z | ratio | p.value |
| --- | --- | --- | --- | --- | --- |
| Fixed effects (OR) | (Intercept) | 0.29 (0.03, 3.28) | -1.01 | 0.29 | 0.315 |
| **Fixed** | **Status** | **3.70 (0.55, 25.05)** | **1.34** | **3.70** | **0.180** |
| Fixed | DayStatus_scale | 2.41 (0.14, 41.33) | 0.61 | 2.41 | 0.544 |
| Fixed | Season.L | 1.29 (0.78, 2.14) | 1.00 | 1.29 | 0.316 |
| Fixed | Season.Q | 1.05 (0.54, 2.03) | 0.14 | 1.05 | 0.886 |
| Fixed | Season.C | 0.75 (0.37, 1.52) | -0.80 | 0.75 | 0.424 |
| Fixed | Status × DayStatus_scale | 5.38 (0.59, 48.87) | 1.50 | 5.38 | 0.135 |
| Random SDs | SD (Intercept) Group | 0.0092 |  |  |  |
| Random SDs | SD (Intercept) MaleID | 0.0611 |  |  |  |
| Random SDs | SD (Day) MaleID | 0.00088 |  |  |  |
| Random correlations | Cor (Intercept ~ Day) MaleID | -0.97 |  |  |  |
| Slopes (Status) | Immigrant | log = 2.56 (-1.30, 6.43) | 1.30 | OR ≈ 12.97 | 0.194 |
| Slopes (Status) | Resident | log = -0.80 (-4.11, 2.50) | -0.48 | OR ≈ 0.45 | 0.634 |
| **Source** | **Term** | **Est (95% CI)** | **z** | **ratio** | **p.value** |
| **Fixed effects (OR)** | (Intercept) | 0.65 (0.04, 10.44) | -0.31 | 0.65 | 0.759 |
| Fixed | Status | 2.30 (0.25, 21.51) | 0.73 | 2.30 | 0.467 |
| Fixed | DayStatus_scale | 9.60 (0.37, 252.17) | 1.36 | 9.60 | 0.175 |
| Fixed | Season.L | 1.89 (1.06, 3.38) | 2.14 | 1.89 | 0.032 |
| Fixed | Season.Q | 1.04 (0.49, 2.21) | 0.10 | 1.04 | 0.920 |
| Fixed | Season.C | 0.71 (0.31, 1.59) | -0.84 | 0.71 | 0.401 |
| Fixed | Status × DayStatus_scale | 4.27 (0.31, 58.26) | 1.09 | 4.27 | 0.276 |
| Random SDs | SD (Intercept) Group | 0.0022 |  |  |  |
| Random SDs | SD (Intercept) MaleID | 0.0231 |  |  |  |
| Random SDs | SD (Day) MaleID | 0.00114 |  |  |  |
| Random correlations | Cor (Intercept ~ Day) MaleID | 0.96 |  |  |  |
| Slopes (Status) | Immigrant | log = 3.71 (-1.05, 8.48) | 1.53 | OR ≈ 41.1 | 0.127 |
| Slopes | Resident | log = 0.81 (-2.70, 4.32) | 0.45 | OR ≈ 2.25 | 0.651 |
| Slope contrast | Immigrant - Resident | log = 2.90 (-2.67, 8.48) | 1.09 | — | 0.276 |
| Contrast | Immigrant - Resident | prob = 0.055 (0.033-0.093) |  | 0.421 | 0.0057 |

**Table S5a:**
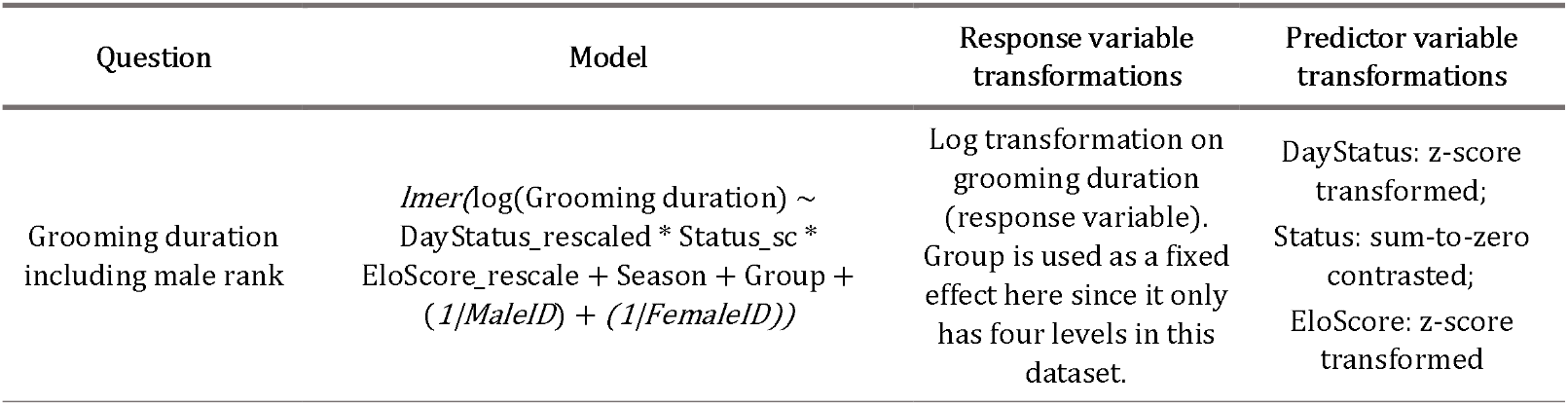
Model used in additional analysis of males including their rank. Note that group identity here is included as a fixed effect since we only had four levels for group in our subset.

**Table S5b,c:** Model output of additional analysis including male rank. Fixed effects are presented as ratios (exponentiated coefficients) with 95% confidence intervals, z-values, and associated *p*-values. Random effect standard deviations (SDs) are reported on the model’s link scale. Estimated marginal means (EMMs) are shown on the response scale, with contrasts expressed as ratios (with 95% CIs) and corresponding test statistics. Slopes of *DayStatus_rescale* are given per +1 SD change, with slope contrasts reported on the link scale.

**b. Female-given grooming durations based on the male rank model.**
| Source | Term | Est (95% CI) | t | ratio | p |
| --- | --- | --- | --- | --- | --- |
| Fixed effects (Ratio) | (Intercept) | 4.584 (3.90, 5.26) | 13.10 | 97.8 | <0.001 |
| Fixed effects (Ratio) | DayStatus_rescale | -0.081 (-0.326, 0.163) | -0.65 | 0.92 | 0.984 |
| Fixed effects (Ratio) | Status (Resident) | 0.117 (-0.139, 0.374) | 0.90 | 1.12 | 0.865 |
| Fixed effects (Ratio) | EloScore_rescale | 0.074 (-0.192, 0.341) | 0.55 | 1.08 | 0.801 |
| Fixed effects (Ratio) | Season.L | -0.028 (-0.445, 0.390) | -0.13 | 0.97 | 0.585 |
| Fixed effects (Ratio) | Season.Q | -0.043 (-0.483, 0.397) | -0.19 | 0.96 | 0.585 |
| Fixed effects (Ratio) | Season.C | -0.233 (-0.739, 0.273) | -0.90 | 0.79 | 0.585 |
| Fixed effects (Ratio) | Group (BD) | 0.253 (-0.454, 0.961) | 0.70 | 1.29 | 0.850 |
| Fixed effects (Ratio) | Group (KB) | 0.072 (-0.602, 0.747) | 0.21 | 1.07 | 0.850 |
| Fixed effects (Ratio) | Group (NH) | 0.214 (-0.492, 0.921) | 0.60 | 1.24 | 0.850 |
| Fixed effects (Ratio) | DayStatus_rescale × Status | -0.173 (-0.380, 0.034) | -1.65 | 0.84 | 0.113 |
| Fixed effects (Ratio) | DayStatus_rescale × EloScore | -0.038 (-0.268, 0.192) | -0.33 | 0.96 | 0.966 |
| Fixed effects (Ratio) | Status × EloScore | 0.066 (-0.209, 0.340) | 0.47 | 1.07 | 0.812 |
| Fixed effects (Ratio) | DayStatus × Status × EloScore | -0.053 (-0.278, 0.172) | -0.46 | 0.95 | 0.648 |
| Random effects | SD (Intercept) FemaleID | 0.000 |  |  |  |
| Random effects | SD (Intercept) MaleID | 0.213 |  |  |  |
| Random effects | SD Residual | 0.951 |  |  |  |
| Slope (Status \| DayStatus) | Immigrant | -0.259 (-0.663, 0.146) | -1.26 | 0.77 (0.52, 1.16) | 0.209 |
| Slope (Status \| DayStatus) | Resident | 0.093 (-0.168, 0.354) | 0.71 | 1.10 (0.85, 1.42) | 0.480 |
| Contrast slopes (Status \| DayStatus) | Immigrant – Resident | -0.352 (-0.780, 0.076) | -1.61 | 0.70 (0.46, 1.08) | 0.110 |
| Slope (Status \| EloScore) | Immigrant | 0.141 (-0.38, 0.66) | 0.53 | 1.15 | 0.595 |
| Slope (Status \| EloScore) | Resident | 0.009 (−0.19, 0.21) | 0.08 | 1.01 | 0.935 |
| Contrast slopes (Status \| EloScore) | Immigrant – Resident | 0.133 (−0.44, 0.71) | 0.459 | 1.14 | 0.647 |

**c. Male-given grooming duration based on the male rank model.**
| Source | Term | Est (95% CI) | t | ratio | p |
| --- | --- | --- | --- | --- | --- |
| Fixed effects (Ratio) | (Intercept) | **3.823 (2.87, 4.78)** | 7.88 | **45.8** | **<0.001** |
| Fixed effects (Ratio) | DayStatus_rescale | **0.195 (−0.127, 0.517)** | 1.19 | **1.21** | **0.604** |
| Fixed effects (Ratio) | Status (Resident) | **−0.478 (−0.880, −0.077)** | −2.34 | **0.62** | **0.017** |
| Fixed effects (Ratio) | EloScore_rescale | **−0.277 (−0.644, 0.091)** | −1.50 | **0.76** | **0.150** |
| Fixed effects (Ratio) | Season.L | **−0.221 (−0.682, 0.240)** | −0.94 | **0.80** | **0.756** |
| Fixed effects (Ratio) | Season.Q | **−0.025 (−0.604, 0.554)** | −0.08 | **0.98** | **0.756** |
| Fixed effects (Ratio) | Season.C | **0.275 (−0.372, 0.922)** | 0.83 | **1.32** | **0.756** |
| Fixed effects (Ratio) | Group (BD) | **0.260 (−0.777, 1.298)** | 0.52 | **1.30** | **0.605** |
| Fixed effects (Ratio) | Group (KB) | **0.589 (−0.400, 1.579)** | 1.17 | **1.80** | **0.605** |
| Fixed effects (Ratio) | Group (NH) | **0.224 (−0.787, 1.235)** | 0.43 | **1.25** | **0.605** |
| Fixed effects (Ratio) | DayStatus × Status | **0.194 (−0.093, 0.481)** | 1.32 | **1.21** | **0.311** |
| Fixed effects (Ratio) | DayStatus × EloScore | **0.127 (−0.163, 0.417)** | 0.86 | **1.14** | **0.726** |
| Fixed effects (Ratio) | Status × EloScore | **−0.108 (−0.471, 0.255)** | −0.58 | **0.90** | **0.996** |
| Fixed effects (Ratio) | DayStatus × Status × EloScore | **0.125 (−0.162, 0.412)** | 0.85 | **1.13** | **0.395** |
| Random effects | SD (Intercept) FemaleID | 0.272 |  |  |  |
| Random effects | SD (Intercept) MaleID | 0.374 |  |  |  |
| Random effects | SD Residual | 0.867 |  |  |  |
| Slope (Status \| DayStatus) | Immigrant | 0.375 (−0.161, 0.912) | 1.37 | 1.46 (0.85, 2.49) | 0.174 |
| Slope (Status \| DayStatus) | Resident | 0.001 (−0.296, 0.298) | 0.01 | 1.00 (0.74, 1.35) | 0.995 |
| Contrast slopes (Status \| DayStatus) | Immigrant – Resident | 0.374 (−0.200, 0.948) | 1.28 | 1.45 (0.82, 2.58) | 0.204 |
| Slope (Status \| EloScore) | Immigrant | −0.380 (−0.99, 0.23) | −1.098 | 0.68 | 0.274 |
| Slope (Status \| EloScore) | Resident | −0.169 (−0.45, 0.11) | −1.203 | 0.84 | 0.235 |
| Contrast slopes (Status \| EloScore) | Immigrant – Resident | −0.211 (−0.95, 0.53) | −0.565 | 0.81 | 0.574 |

